# Caffeine Impairs Red Blood Cell Storage Quality by Dual Inhibition of ADORA2b Signaling and G6PD Activity

**DOI:** 10.1101/2025.05.27.656446

**Authors:** Monika Dzieciatkowska, Ariel Hay, Aaron Issaian, Gregory R. Keele, Shaun Bevers, Travis Nemkov, Julie A Reisz, Mark Maslanka, Daniel Stephenson, Amy L Moore, Xutao Deng, Mars Stone, Kirk C. Hansen, Steve Kleinman, Philip J Norris, Michael P. Busch, Grier P Page, Nareg Roubinian, Yang Xia, James C Zimring, Angelo D’Alessandro

## Abstract

Caffeine is the most widely consumed psychoactive substance globally, yet its peripheral physiological effects remain incompletely understood. Leveraging comprehensive data from 13,091 blood donors in the REDS RBC-Omics study, we identify caffeine as a significant modulator of red blood cell (RBC) storage quality and transfusion outcomes. Elevated caffeine levels were reproducible across multiple donations from 643 recalled donors, selected based on their extremes in hemolytic propensity. Both in the screening and recalled cohorts, higher caffeine levels were associated with disrupted RBC metabolism, characterized by reduced glycolysis, depletion of adenylate pools or 2,3-bisphosphoglycerate, and increased markers of oxidative stress and osmotic fragility, including kynurenine accumulation. These observations were recapitulated in plasma and RBCs of eight volunteers upon consumption of a cup of coffee independently of brewing method (Chemex vs espresso). Clinically, elevated caffeine correlated with increased hemolysis and lower post-transfusion hemoglobin increments, especially pronounced in recipients transfused with RBCs from donors carrying common polymorphisms in the ADORA2b gene, a key regulator of RBC metabolism in hypoxia. These human findings were mechanistically validated using a murine model deficient in ADORA2b, which demonstrated impaired glycolytic flux, compromised antioxidant defenses – including caffeine-dependent direct inhibition of recombinantly-expressed glucose 6-phosphate dehydrogenase, and decreased transfusion efficacy (lower hemoglobin increments, higher bilirubin post-transfusion), effects further exacerbated by caffeine exposure during storage. Our study positions caffeine consumption as a modifiable factor in blood transfusion practice, advocating for precision strategies that integrate genetic and exposome factors, and identifies metabolic interventions to enhance blood quality and clinical outcomes.

**One sentence summary:** Caffeine consumption and genetic variants in the ADORA2b receptor synergistically impair red blood cell metabolism and transfusion efficacy, revealing a modifiable exposome–gene interaction for precision transfusion medicine.

## INTRODUCTION

Caffeine is one of the most widely consumed psychoactive compounds worldwide, primarily sourced from coffee, tea, energy drinks, and sodas (1). In the United States, over 67% of adults report daily coffee consumption, averaging more than 1.5 cups per day (∼135 mg), with 36% of adults reporting an average daily consumption of 3-5 cups. Globally, coffee consumption exceeds 10 billion kilograms annually and continues to grow at an estimated 5% per year (2). An 8-ounce cup of brewed coffee (∼240 ml) contains approximately 95 mg of caffeine, while a 12-ounce can of cola (∼355 ml) contains around 35 mg, and a standard energy drink may contain up to 160 mg. These widespread dietary exposures make caffeine one of the most pervasive modulators of systemic and cellular physiology.

As a non-selective antagonist of adenosine receptors, caffeine – a purine alkaloid - exerts its primary effects through blockade of A1 and A2A receptors in the central nervous system, enhancing alertness and psychostimulation (3). However, peripheral adenosine receptors — notably the A2B subtype (ADORA2B) — are also affected. Mature red blood cells (RBCs), though anucleate, express functional ADORA2B (4), a G protein-coupled receptor that activates cAMP production and downstream kinases such as protein kinase A (PKA) and AMP-activated protein kinase (AMPK) (5). In hypoxic conditions, adenosine levels increase, activating ADORA2B to stimulate glycolysis and 2,3-bisphosphoglycerate (2,3-BPG) synthesis (4). This facilitates oxygen unloading at the tissue level (6) and supports redox balance by favoring fluxes through glycolysis at the expense of the pentose phosphate pathway (PPP) (4), thereby liming NADPH generation and thus NADPH-dependent glutathione recycling.

The clinical relevance of these pathways becomes evident in the context of blood storage, when oxidant stress to RBCs is high (7). Transfusion of packed RBCs is the most common inpatient procedure after vaccination in the U.S., with over 12 million units transfused annually. RBCs are stored for up to 42 days at 1–6°C, during which they undergo progressive biochemical deterioration (7). This “storage lesion” includes depletion of ATP and 2,3-BPG, oxidative damage to proteins and lipids, vesiculation, and membrane loss (7). Importantly, the capacity of stored RBCs to regenerate 2,3-BPG and maintain ATP upon rewarming is tightly linked to their capacity to circulate in vivo after transfusion, a gold standard parameter that assesses the quality of transfused blood, also referred to as post-transfusion recovery (PTR) (8).

Stored RBCs progressively lose the capacity to activate the PPP (9), resulting in decreased NADPH production and loss of antioxidant buffering capacity. This makes RBCs vulnerable to oxidative hemolysis (10), particularly in recipients with inflammatory conditions. Accumulation of purine catabolites such as hypoxanthine, a marker of oxidant stress and ATP degradation, further correlates with reduced RBC lifespan after transfusion (8). Therefore, pathways that preserve purine homeostasis and redox balance are critical targets for improving stored RBC quality.

High-altitude physiology offers a natural model for understanding the role of purine metabolism in RBC energy and redox physiology (11). At elevations above 3,000 meters, plasma adenosine levels rise by up to 10-fold, activating ADORA2B in RBCs to increase 2,3-BPG and enhance tissue oxygen delivery (4). In humans and mice, this adaptation is blunted when adenosine is enzymatically degraded (12) or ADORA2B is genetically deleted. In murine models, pharmacologic activation of AMPK improves post-storage 2,3-BPG and ATP levels(13). These mechanisms are beneficial to acclimatization to high altitude, to the extent that proteasomal degradation of the adenosine equilibrative nucleoside transporter ENT1 favors faster acclimatization upon reascent (14). Caffeine, by antagonizing ADORA2B, has the potential to impair this protective signaling.

Additionally, caffeine has been reported to directly and competitively bind to G6PD— the rate-limiting enzyme of the PPP—inhibiting its activity and reducing NADPH production (15). Xu and colleagues report that, in vitro, caffeine - at concentrations as low as 50 μM (equivalent to 1–2 cups of coffee) - reduces G6PD activity by up to 40%. Thus, habitual caffeine consumption could plausibly affect the redox capacity and glycolytic balance of donor RBCs, particularly during storage.

Beyond biological (age, BMI, sex) or genetic factors (16), the quality of stored blood is impacted by the so-called exposome, i.e., donors’ dietary, recreational, professional or medical exposures that are not grounds for blood donor deferral (17). Two otherwise healthy blood donors may produce RBC units of vastly different storage quality due to recent dietary intake, medications, or occupational exposures.

Encouraged by interesting preliminary data (18), here we investigated the impact of caffeine on RBC storage quality, here we stratified 13,000 donors enrolled in the Recipient Epidemiology and Donor Evaluation Study (REDS) RBC Omics, with detailed genomic, metabolomic, and storage phenotype data. Leveraging this resource, we conducted a two-part investigation to determine how caffeine and ADORA2B signaling interact to shape RBC storage outcomes, including in vitro and post-transfusion hemolysis.

## Materials and Methods

In the interest of space, extensive details for this section are provided in **Supplementary Materials.**

### REDS RBC Omics: Index and Recalled donors

Metabolomics analyses were performed on day 42 packed RBCs from 13,091 “index” donors enrolled across four blood centers as part of the Recipient Epidemiology and Donor evaluation Study (REDS RBC Omics). Omics results were analyzed as a function of single nucleotide polymorphisms (SNPs) mapping on the region coding for ADORA2b, as extrapolated from genomics data on 879,000 single nucleotide polymorphisms (SNPs). Donors ranking in the 5^th^ and 95^th^ percentile for end-of-storage hemolysis (n=643) were invited to donate a second (“recalled”) unit, which was tested at storage days 10, 23 and 42 (1,929 samples) for hemolytic parameters, high-throughput metabolomics (19–21), proteomics (22), lipidomics (23). Results were analyzed as a function of caffeine levels and ADORA2b SNPs.

### Caffeine consumption in healthy volunteers

To interrogate the metabolic effect of caffeine consumption we randomized eight volunteer to consume a coffee cup either brewed as chemex or espresso (n=4 per group). Blood samples were collected at baseline and after 45 min and 5h from consuming the beverage, and plasma and RBCs were separated via gentle centrifugation (10 min at 4°C at 2,000 *g*) prior to metabolomics analyses.

### G6PD activity assay in presence of caffeine

We recombinantly expressed and purified a human canonical G6PD or the deficient African variant (V68M; N126D – class III variant, 10-60% residual activity) – most prevalent in the REDS RBC Omics cohort (∼13% of donors of African descent and 2% of donors of Hispanic descent (24)). Activity assays were performed as described (15), in presence or absence of 50 μM caffeine (15).

### Storage and Post-transfusion recovery (PTR) studies in ADORA2b mice

Mouse post-transfusion recovery (PTR) studies were performed as previously described (25). Storage of ADORA2b KO mice (n=3) for 7 or 12 days was followed by transfusion into Ubi-GFP mice, which were used as recipients to allow visualization of the test cells in the non-fluorescent gate. To control for differences in transfusion and phlebotomy, mCherry (red fluorescent-labeled) RBCs were used as a tracer RBC population (never stored). These RBCs were added to stored RBCs immediately prior to transfusion. PTR was calculated by dividing the post-transfusion by the pre-transfusion ratio (Test/Tracer), with the maximum value set as 1 (or 100% PTR). In follow up experiments, PTR studies were repeated after supplementation of caffeine (100 μM – no. C0750 Sigma Aldrich) to citrate-phosphate-dextrose-adenine CPDA1 standard storage solutions (25). This concentration is consistent with physiological plasma caffeine levels in high-consumption donors (low to high range: 20–100 μM (18)) and in vitro studies mimicking real-life exposures (range 50–100 μM (15)).

### Determination of hemoglobin and bilirubin increment via the vein-to-vein database

Association of ADORA2b alleles with hemoglobin increments was performed by interrogating the vein-to-vein database, as described in Roubinian et al.(26). Analyses were performed on the whole REDS population and further stratified for caffeine measurements, with a focus on the upper 50^th^ percentile of caffeine measurements in REDS donors.

## RESULTS

### High levels of caffeine in donor red blood cells are reproducible across time

To assess the extent and physiological impact of caffeine exposure on stored red blood cell (RBC) quality, we first quantified caffeine levels in packed RBC units from 13,091 donors enrolled in the REDS RBC Omics study. Caffeine levels were either below detection threshold or negligible in ∼5,000 donors, spanning a broad and continuous distribution (**Figure 1A**). In the full cohort, caffeine levels correlated to metabolic changes in metabolism, including a negative association with branched chain amino acids (Valine, Isoleucine, Threonine) and the related carnitine-conjugated ketoacid (AC 5-OH), with positive association with the bacterial metabolites 3-sulfocatechol and Hippurate, kynurenine metabolites (8-methoxykynurenate), and lipid oxidation products (prostaglanding A2/B2 isomers, several acyl-carnitines (AC – **Figure 1B**).

**Fig. 1.**
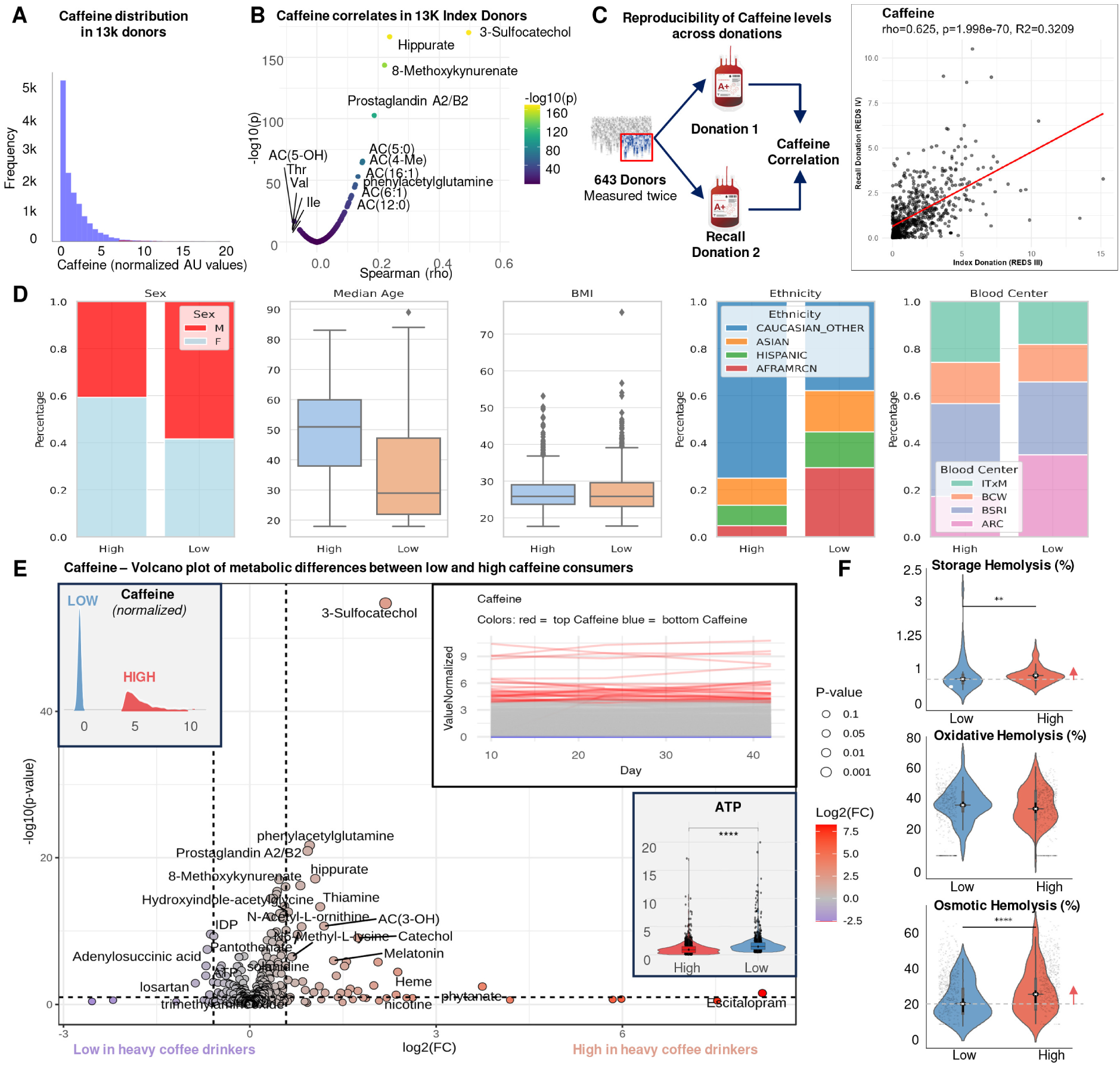
– Caffeine, donor biology and red blood cell (RBC) hemolysis in 13,091 REDS RBC Omics donors. Caffeine distribution in 13,091 packed RBC units from donors enrolled in the REDS RBC Omics study (**A**). In **B**, metabolic correlates to caffeine levels in the same blood units. In **C**, caffeine levels were reproducible across multiple independent donations 6-12 months apart from 643 Index donors who were also enrolled in the Recalled phase of the study, owing to their extreme hemolytic propensity (<5^th^ or >95^th^ percentile). In **D**, stratification of REDS Index donors based on caffeine levels (top and bottom 1,000 donors by high or low caffeine levels) as a function of donor sex, age, body mass index (BMI), self-reported ethnicity and blood center in which the donors enrolled. In **E**, the volcano plot highlight the significant metabolic changes between caffeine low and high donors (n=1,000 per group – ATP is highlighted in the violin plot in the bottom right corner) in REDS Index donors at storage day 42. In the recalled donor cohort (n=643), caffeine levels did not change in packed RBCs when dosed at storage day 10, 23 and 42 (indent line plot). In **F**, blood units with high caffeine levels were associated with significantly higher storage ad osmotic hemolysis (** p<0.01; **** p<0.0001).

For a subset of donors (extreme hemolyzers) enrolled both in the Index and Recalled phase of the REDS RBC Omics study (n=643), caffeine levels in packed RBCs were reproducible across multiple donations 6–12 months apart (**Figure 1C**). Stratification of donors based on high vs low caffeine levels (n=1,000 per group) revealed interesting demographic trends, with higher levels dosed in older individuals, females under 51, and donors with lower BMI, especially donors of Caucasian descent enrolled by the American Red Cross (**Figure 1D**). Similar trends were observed for the primary caffeine metabolite, paraxanthine (80% of caffeine is metabolized to paraxanthine), which was higher in older donors or younger females of Caucasian descent, showing a strong positive correlation (0.7 Spearman) and stable ratios across donor ages and sex (**Supplementary Figure 1A-C**). However, paraxanthine to caffeine ratios, a proxy for caffeine metabolism, were tendentially higher in younger donors, especially females of African American, Asian and Hispanic descent (**Supplementary Figure 1A-D**). Metabolite quantitative trait loci (mQTL) analysis identified a region on chromosome 19 coding for cytochrome p450 2A6 (CYP2A6) associated with paraxanthine and paraxanthine/caffeine levels in the REDS Index cohort (**Supplementary Figure 2A-B**). Top SNPs included rs7251570, rs76112798 and rs11667314, which were most prevalent in donors of Asian or European descent and significantly influenced paraxanthine metabolism (**Supplementary Figure 2C-D**).). Similar findings were recapitulated for the second most abundant caffeine metabolite (12% of total), theobromine, also mapping on the CYP2A6 region (**Supplementary Figure 3A-B**).

### Caffeine levels in REDS donors or coffee consumption in volunteers is associated with altered metabolism and increased hemolysis

Comparative metabolomics between donors with the highest and lowest RBC caffeine levels identified significant alterations in central metabolism, including reduced glycolytic intermediates, increased purine catabolites, and changes in amino acid and redox pathways (**Figure 1.E**).

To isolate the impact of caffeine on the circulating metabolome, we performed metabolomics analyses on plasma and RBCs from eight volunteers after consuming a cup of coffee (chemex or espresso – **Supplementary Figure 4**). Results showed transient elevation of circulating caffeine within 45 min, and rapid metabolism by 300 min. Elevation in sulfocatechol and other bacterial metabolites was observed in plasma and RBCs, consistent with caffeine’s impact on the gut microbiome. Caffeine consumption was associated with a drop in RBC 2,3-BPG and an increase in markers of membrane lipid oxidation and remodeling, including free fatty acids, (hydroxy)acyl-carnitines and lipid hydroperoxides (20:4 HPETE – **Supplementary Figure 4**). Elevation of plasma 2,3-BPG over time suggested release from RBCs, possibly via minor hemolysis.

In the Recalled cohort (n=643), for which samples were collected at storage days 10, 23, and 42, caffeine levels remained stable (**Figure 1E-inset**), indicating that associated metabolic changes were not due to caffeine degradation. High caffeine levels correlated with significantly lower ATP and pentose phosphate pathway intermediates, suggesting impaired energy/redox metabolism and increased hemolytic propensity. Indeed, higher spontaneous and osmotic hemolysis at day 42 were observed in the high caffeine group (p<0.01 and p<0.0001, respectively – **Figure 1F**).

### RBCs with high caffeine show redox imbalance, poor metabolic fitness, and impaired post-transfusion efficacy

Metabolomics, proteomics, and lipidomics analyses of RBCs from Recalled donors (n=100) revealed subtle but consistent separation by caffeine levels in 3D uMAP projections (**Figure 2A–B**). High caffeine levels tracked with increased donor age and were enriched in women under 51 (**Figure 2C**). Longitudinal metabolomics revealed sustained increases in hypoxanthine and kynurenine (**Figure 2D–E**), markers of oxidative stress previously associated with poor PTR. Higher osmotic fragility was confirmed (**Figure 2F**). Hive plot correlation analyses revealed altered metabolic networks in high caffeine units, with weaker associations between ATP or 2,3-BPG and hemolysis (**Figure 2G**), metabolites previously linked to better PTR and oxygen off-loading (8, 27–30).

**Fig. 2.**
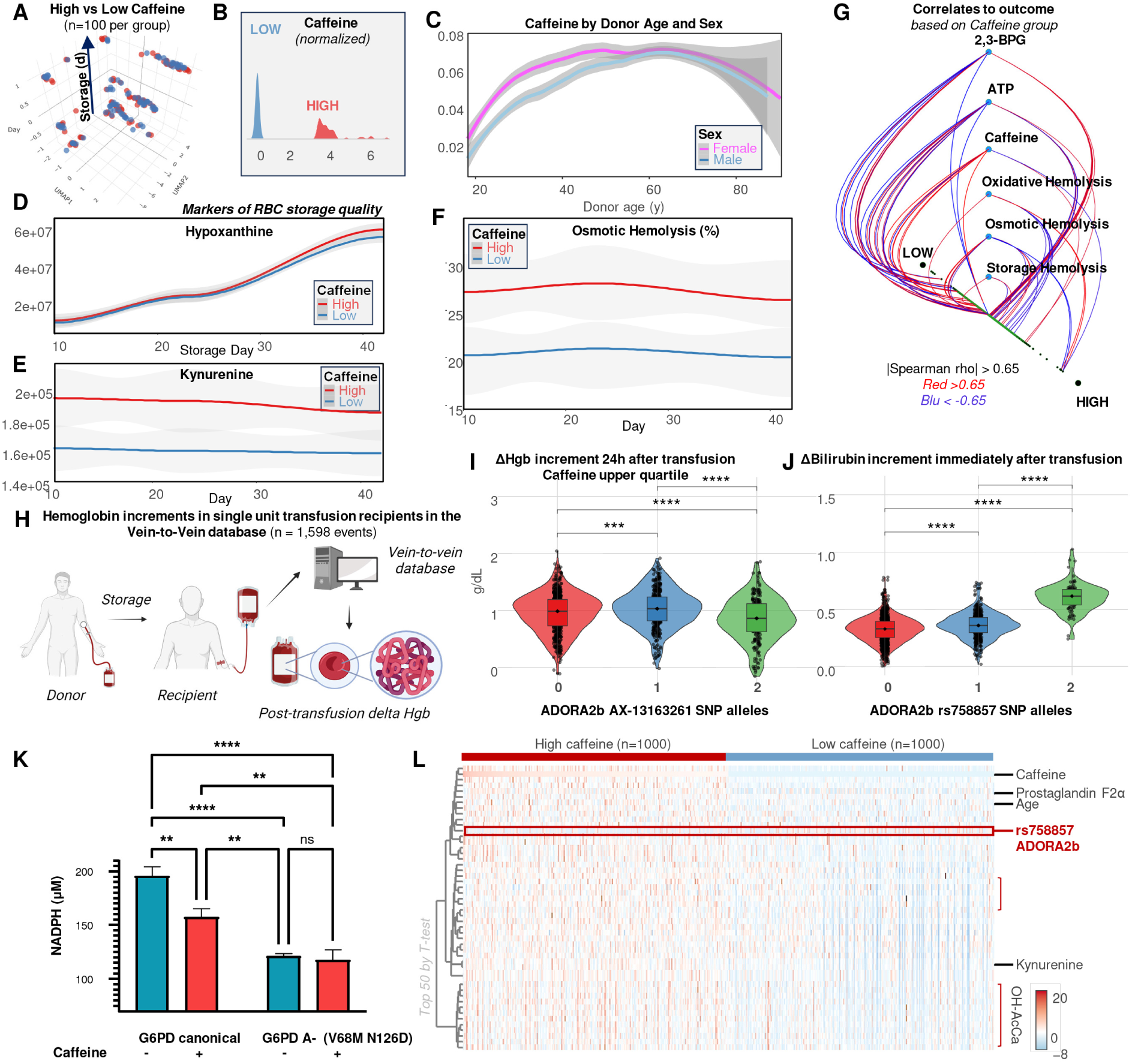
– Higher caffeine levels are associated with elevated osmotic fragility of 643 recalled donors packed RBC and lower hemoglobin increments in transfusion recipients. The 3D uMAP in **A** shows substantial overlap between packed RBCs with high or low caffeine (n=100 per group – distributions in **B**) as a function of storage duration (z axis). Caffeine was higher in donors older than 30, while higher caffeine levels were observed in female donors below the age of 51 compared to males (**C**). In **D-E**, blood units with elevated caffeine show significantly higher levels of metabolic markers of poor post-transfusion recovery, hypoxanthine (p<0.05 at storage day 42) and kynurenine (p<0.01 through the whole storage period). Recalled donor blood units with high caffeine showed significantly higher levels of osmotic hemolysis (**F**). In **G**, hive plot summarizes changes in correlation networks of low and high caffeine blood units to hemolysis parameters (spontaneous storage hemolysis, or after osmotic or oxidant stress), and to functional metabolic parameters, including adenosine triphosphate (ATP) and 2,3-bisphosphoglycerate (2,3-BPG). Upomn interrogation of the REDS vein-to-vein database (**H**), transfusion of packed RBCs with high caffeine (or from donors repeatedly identified as heavy caffeine consumers – highest quartile) results in lower hemoglobin increments after transfusion, especially when the donors are also carriers of one or two alternative alleles for the single nucleotide polymorphism (SNP) for the adenosine receptor ADORA2b AX-13163261 (**I**). Similarly, transfusion of units from donors carrying 1 or 2 alleles for the ADORA2b rs758857 SNP was associated with higher delta-bilirubin increments, independently of caffeine levels (**J**). In vitro incubation of recombinantly expressed canonical or African G6PD variant (V68M; N126D) with 50 µM caffeine inhibited canonical but not deficient G6PD activity (**K**). In **L**, heat map of the top 50 significant differences between high and low caffeine level blood units in REDS RBC omics Index donors identifies donor age, markers of osmotic fragility (kynurenine), markers of lipid peroxidation (hydroxyacyl-carnitines and prostaglandin F2α) and alternative alleles for ADORA2b SNP rs758857 as the most significant differences between the two groups (high and low caffeine – n=1,000).

Transfusion data from the REDS vein-to-vein database (**Figure 2H**) showed significantly lower hemoglobin increments in recipients of high-caffeine RBC units, particularly among donors carrying the ADORA2B SNP AX-13163261 (**Figure 2I**). The ADORA2B SNP rs758857 was also associated with higher increases in bilirubin post-transfusion, a hemolysis marker, independent of donor age or pre-transfusion bilirubin levels (n=1,598 - **Figure 2J, Supplementary Figure 5**).

In vitro, 50 µM caffeine inhibited recombinant G6PD activity but did not further decrease the activity of the African variant (V68M; N126D), suggesting partial inhibition may explain metabolic defects (**Figure 2K**). Integrative omics and genomic analyses highlighted differences in membrane stability and ADORA2B variant prevalence among the top 50 discriminants between high and low caffeine donors (**Figure 2L**).

### ADORA2b genetic variants are prevalent and associate with increased RBC hemolysis

We analyzed 879,000 SNPs in the REDS cohort focusing on ADORA2B. Several variants were common, including AX-13163261 (46.9% heterozygous, 19.9% homozygous – **Figure 3A–B**). Variants rs758857 and AX-13163261 were strongly associated with osmotic (p<e-50) and oxidative hemolysis (p<e-9 -**Figure 3C**), though not storage hemolysis. These alleles were enriched in older donors (rs72821748), those with higher BMI (rs758857), and African Americans (**Figure 3D–F**, **Supplementary Figure 6**). Metabolomics linked these SNPs to altered redox metabolism and elevated caffeine levels (**Figure 3G–I**).

**Fig. 3.**
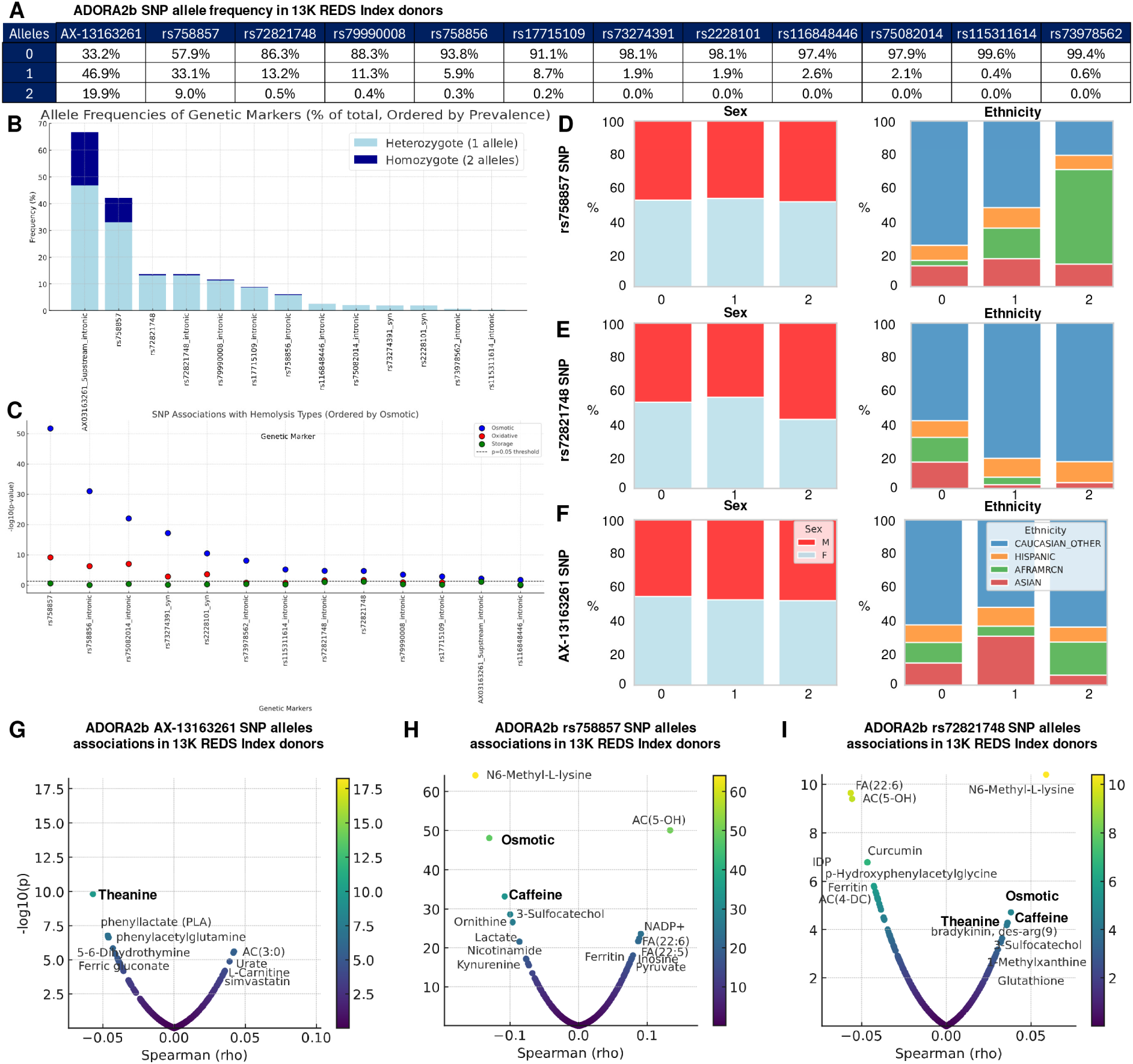
– Prevalence of ADORA2b SNPs in REDS Index donors and association with hemolytic propensity. A list of the ADORA2b SNPs assayed in the 13,091 REDS Index donors is summarized in **A**, and the prevalence of heterozygosity or homozygosity for the alternative alleles is show in **B**. In **C**, SNP association to osmotic (blue), oxidative (red) or storage hemolysis (green). In **D-F**, stratification of the ADORA2b SNPs most strongly associated with hemolysis (especially osmotic and oxidative) by donor demographics (sex and ethnicity; age and demographics are shown in **Supplementary Figure 1**). In **G-I**, volcano plots of Spearman correlation of allele copies to metabolites and hemolysis parameters in the same SNPs.

### ADORA2b knockout impairs glycolysis and redox homeostasis in stored murine RBCs

To model mechanisms, we stored RBCs from WT and ADORA2B KO mice for 0, 7, or 12 days (equivalent to human 0, 21, 42 days – **Figure 4A-B**). Untargeted metabolomics showed separation by genotype and storage duration (**Figure 4C**). Combined with tracing data from 1,2,3-^13^C_3_-glucose experiments, our results confirmed impaired glycolytic flux in KO RBCs (**Figure 4D**), with reductions in intermediates of glycolysis, PPP, and glutathione metabolism (**Figure 4E, Figure 5**). KO RBCs had lower adenylate pools (calculated as ATP+0.5 ADP/(ATP+ADP+AMP)) and altered GSH/GSSG ratios, consistent with oxidative stress seen in high-caffeine RBCs.

**Fig. 4.**
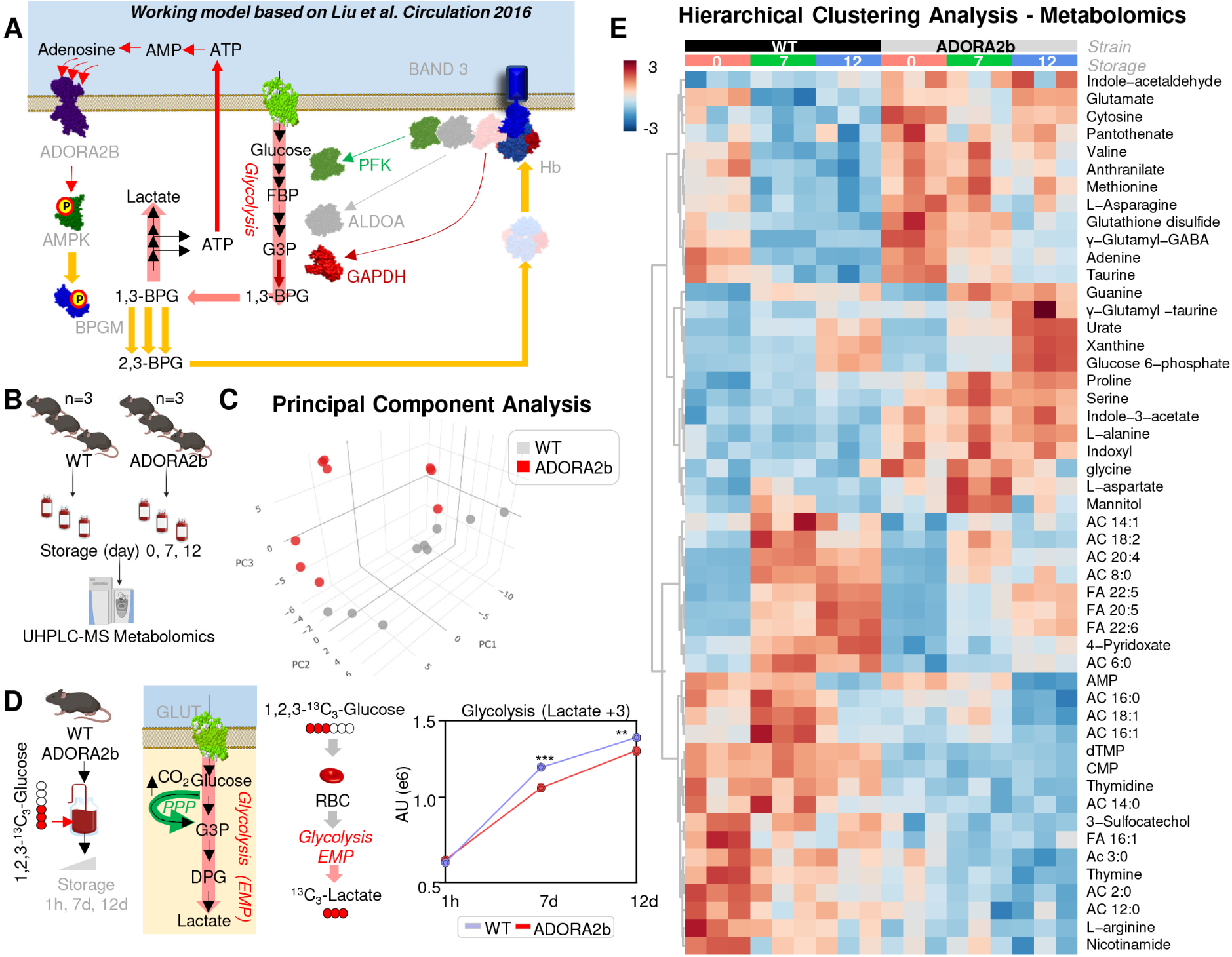
– ADORA2b regulates glycolytic fluxes in stored RBCs. A literature-based summary model of the role of ADORA2b in regulating RBC metabolic fluxes through glycolysis based on Liu et al. Circulation 2016 (**A**). In **B**, packed RBCs from WT C57BL6/J mice (n=3) and ADORA2b KO mice (n=3) were stored for 0, 7, 12 days (equivalent to human RBC storage for 42 days) under refrigerated conditions in standard storage solutions, prior to metabolomics analysis. In **C**, principal component analysis of metabolomics data from this study shows a significant effect of storage duration and genotype. In **D**, the experiment was repeated by supplementing CPDA1 with 5 mM 1,2,3-^13^C_3_-glucose, which affords the direct measurements of metabolic fluxes through glycolysis by measuring the ^13^C_3_-lactate isotopologue. Results indicate a significant decrease in ^13^C_3_-lactate in ADORA2b KO mice. In **E**, heat map of the top 50 metabolites by time series two-way ANOVA (by genotype and storage) in WT vs ADORA2b KO mice at storage day 0, 7 and 12.

**Fig. 5.**
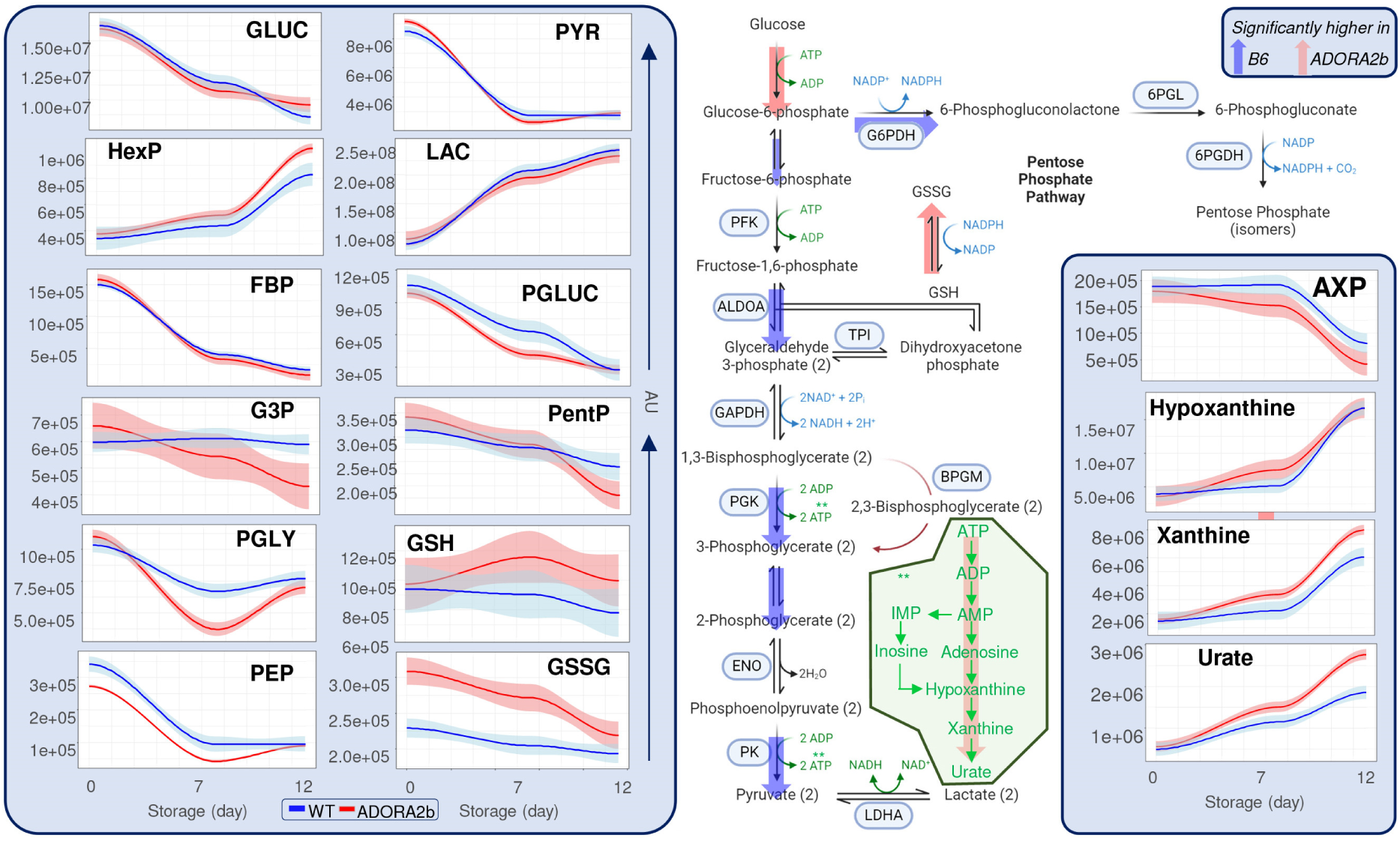
– Overview of glycolysis, the pentose phosphate pathway, glutathione and purine metabolism in stored WT and ADORA2b KO mice. Red lines: ADORA2b; Blue lines: WT; n=3 per group, median ± ranges (solid line and faded band) are shown for each metabolite. Abbreviations: GLUC: glucose; HexP: hexose phosphate (isomers); FBP: fructose bisphosphate; G3P: glyceraldehyde 3-phosphate; PGLY: phosphoglycerate (isomers); PEP: phosphoenol-pyruvate; PYR: pyruvate; LAC: lactate; PGLUC: 6-phosphogluconage; PentP: pentose phosphate (isomers); GSH and GSSG: reduced and oxidized glutathione; AXP: adenylate pool (ATP + 0.5 ADP / (ATP+ADP+AMP).

### Loss of ADORA2b alters redox-sensitive proteins and proteostasis in stored RBCs

We profiled the RBC proteome of WT and KO mice across storage (days 0, 7, 12 – **Figure 6A**). PCA and LDA confirmed genotype-dependent differences (**Figure 6B–C**). Time-series ANOVA identified changes in redox regulators (Psmd5, Ube2l3), glycolysis enzymes (Pkm), cytoskeletal proteins (Tubb4a, Dmtn, Flna), and vesiculation markers (Cltc – **Figure 6D**). Redox proteomics revealed increasing oxidation over time in KO RBCs (**Figure 6E**). Pathway analysis implicated dysregulated proteostasis and antioxidant defenses (**Figure 6F**).

**Fig. 6.**
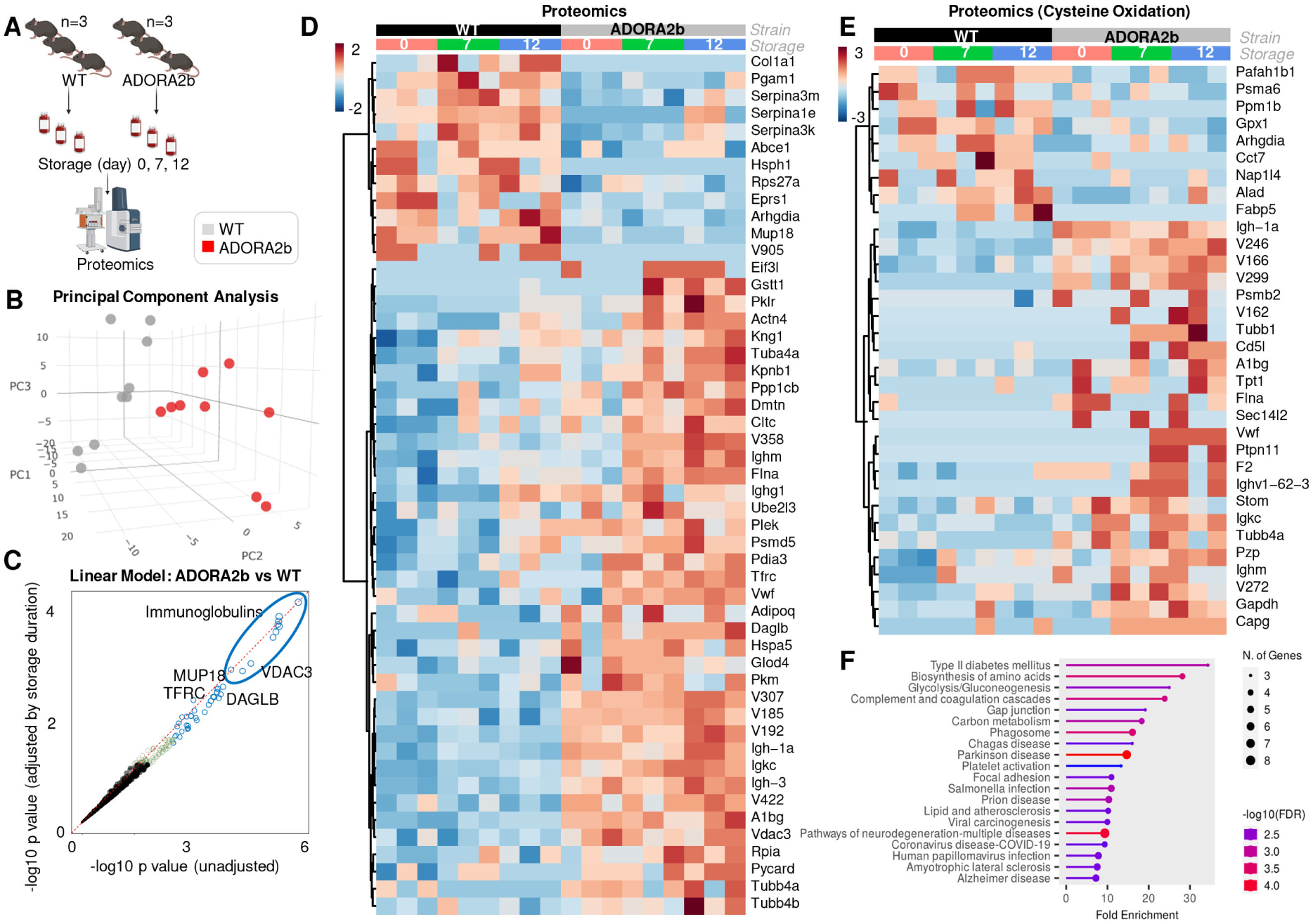
– Proteomics of stored RBCs from WT and ADORA2b knockout. In **A**, overview of the experimental design. Principal component analysis (**B**) and linear discriminant analysis (**C**) of proteomics results based on genotype and storage duration (0, 7 or 12 days). In **D**, heat map of the top 50 significant proteins (official gene symbols) by two-way time series ANOVA as a function of ADORA2b KO and storage duration. In **E**, top 25 sulfenic, sulfonic or cysteine to dehydroalanine redox modifications between WT and ADORA2b KO RBCs during storage. In **F**, pathway analysis based on proteomics results.

### ADORA2b deletion impairs PTR in stored RBCs, an effect exacerbated by caffeine

PTR studies in GFP+ recipient mice transfused with WT or KO RBCs stored 7 or 12 days showed reduced recovery in KO RBCs (**Figure 7A–B**). PTR positively correlated with ATP and glycolytic intermediates (**Figure 7C–D**). Storage with 100 µM caffeine further reduced PTR in KO but not WT RBCs (**Figure 7F**), demonstrating a gene–environment interaction. Clustering of top caffeine- and storage-sensitive metabolites emphasized disruptions in redox and nucleotide metabolism in KO RBCs (**Figure 7E**). A mechanistic model is presented in **Figure 7G**.

**Fig. 7.**
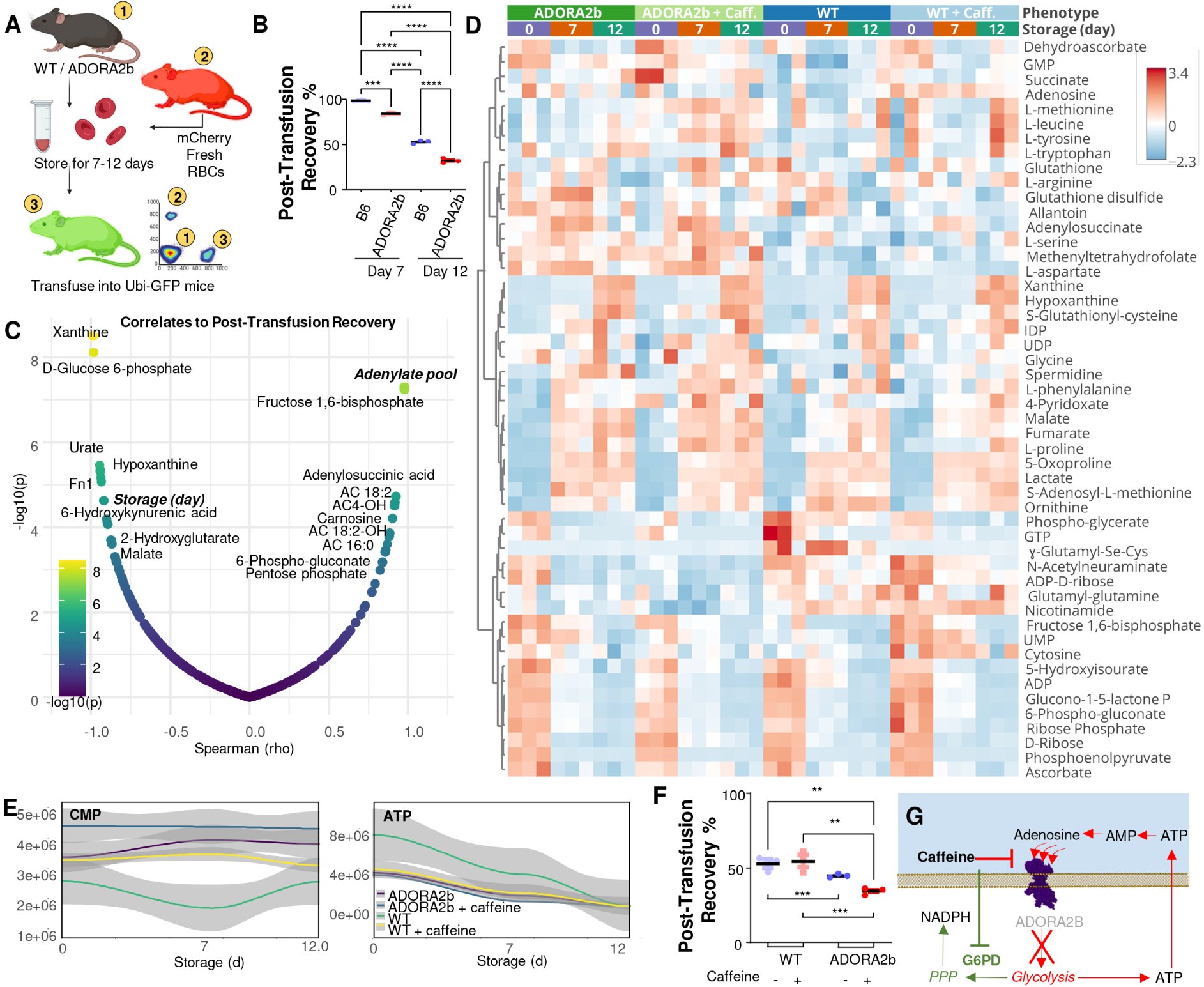
– ADORA2b KO negatively impacts post-transfusion recovery (PTR) of stored RBCs and is aggravated by caffeine supplementation. PTR studies were performed on stored (day 7 and 12) RBCs from WT and ADORA2b KO mice through transfusion into Ubi-GFP mice and co-transfusion with test mCherry fresh WT RBCs (**A**). Results indicate a significant drop in PTR in KO mice (**B**). In **C**, metabolic correlates to PTR. Supplementation of 100 µM caffeine to stored RBCs from WT and ADORA2b KO mice significantly impacted RBC metabolism (top 50 metabolites by two-way ANOVA are shown in the heat map in **D**). Highlights include cytidine monophosphate (CMP) and ATP (**E**). Caffeine supplementation further decreased PTR in ADORA2b KO mice (**F**). A working model is proposed in **G**, based on experimental data and existing literature on the role of caffeine as an ADORA2b antagonist and G6PD inhibitor.

### Adenosine supplementation only partially restored metabolic defects in human stored RBCs

Given the role of adenosine signaling, we tested adenosine supplementation (1, 5, 10 µM) in stored human RBCs (**Supplementary Figure 7**). Adenosine increased intracellular levels early in storage but had modest metabolic impact. Lactate was reduced; ATP, ADP, and 2,3-BPG were higher in the first week. After three weeks, AMP, IMP, and inosine increased in supplemented units, consistent with adenosine catabolism and limited long-term efficacy (**Supplementary Figure 7D–E**).

## DISCUSSION

In the present study we report that caffeine, a ubiquitous dietary component, is a previously underappreciated factor influencing the quality of stored RBCs. By leveraging a large-scale cohort of more than 13,000 blood donors from the REDS RBC-Omics study, we demonstrate that caffeine levels – detected in blood products at biologically relevant thresholds in the micromolar to hundreds of micromolar range (18) - correlate strongly with impaired RBC metabolic integrity and increased storage hemolysis, translating into decreased transfusion efficacy. Stratification of donors by circulating caffeine levels revealed demographic patterns consistent with established population-level consumption trends. Epidemiological surveys in the United States and Europe report higher average caffeine intake among older adults, largely attributable to habitual coffee consumption, which increases with age and plateaus in the 60s (31). Although men typically consume slightly more caffeine overall, studies suggest that women—particularly those under 51—may exhibit higher circulating caffeine concentrations due to slower metabolism influenced by hormonal factors, including estrogen and oral contraceptive use (31). Our measurements of paraxanthine/caffeine ratios are consistent with this model. Caffeine levels also tend to be higher in individuals with lower BMI, likely reflecting a smaller volume of distribution and possibly differences in metabolic clearance. Ethnic variation in both consumption habits and metabolism contributes further: non-Hispanic White individuals report the highest coffee intake in national surveys, a pattern reflected in our cohort, while inter-individual variation in CYP1A2 genotype and expression may additionally shape plasma caffeine levels (32). Collectively, these findings support the interpretation that lifestyle, demographic, and pharmacogenetic factors converge to influence measurable caffeine exposure in blood donors.

While caffeine effects on the central nervous system, cardiovascular disease and cancer (33) are well documented, peripheral actions—especially on mature erythrocytes— have remained underexplored. Here, we show that elevated caffeine in RBC units is consistently associated with a metabolic phenotype indicative of compromised glycolytic activity, reduced ATP, total adenylate pools and 2,3-BPG, as well as with heightened oxidative stress, ferroptosis-like processes (34) and osmotic fragility markers, including oxidized glutathione (35), lipid peroxidation products (34) and hydroxyacyl-carnitines (36), hypoxanthine (8) and kynurenine (28). These findings were validated in volunteers within 45 min to 5h after consuming coffee, independently from brewing methods (espresso vs chemex). These biochemical alterations culminate in increased spontaneous and stress-induced hemolysis during storage, consistent with the increased susceptibility to extravascular hemolysis via splenic sequestration and erythrophagocytosis (37) of storage-damaged RBCs that become energetically effete (38) and accumulate oxidant damage, leading to elevated proteasomal degradation (39, 40). Clinically relevant, recipients of RBC units from donors with high caffeine consumption experienced significantly lower post-transfusion hemoglobin increments. Importantly, these effects were more pronounced in donors harboring specific polymorphisms in the ADORA2b gene, a receptor expressed on RBC membranes for which caffeine is a competitive antagonist, illustrating a critical gene-exposome interaction (17).

Observations in human RBCs are mechanistically supported by our parallel studies in a murine model deficient in ADORA2b (41). While adenosine signaling via ADORA2b signaling has been previously linked to acclimatization to high altitude hypoxia by favoring energy metabolism to boost oxygen off-loading (4), our findings here underscore the critical role of purinergic signaling in maintaining RBC metabolism during refrigerated storage. Inhibition or genetic deletion of ADORA2b disrupts this metabolic adaptation, mirroring the metabolic dysregulation observed with caffeine exposure. In line with this, ADORA2b knockout mice exhibited markedly diminished glycolytic flux, reduced ATP levels, and impaired glutathione recycling during RBC storage, phenocopying the human high-caffeine RBC phenotype. The subsequent decline in RBC viability post-transfusion in knockout mice was further exacerbated by caffeine supplementation during storage, confirming the synergistic detriment of ADORA2b inhibition and caffeine exposure.

Notably, our results confirms and expands upon previous hypotheses implicating caffeine as a direct inhibitor of G6PD (15). Our direct enzymatic assays reveal inhibitory effects of caffeine on canonical G6PD activity at physiologically relevant concentrations. However, the effect did not further compound with G6PD deficiency in the African variant. Therefore, the adverse effects of caffeine observed in stored RBC units are potentially also direct, mediated both through disrupted purinergic signaling (ADORA2b) and direct enzymatic inhibition (G6PD), in part mimicking the increased susceptibility to hemolysis and impaired post-transfusion survival of RBCs from G6PD deficient donors(24, 26, 35, 42).

The translational implications of our findings are significant. First, donor caffeine consumption—a common dietary exposure for up to 75% of Americans—emerges as a modifiable behavioral factor potentially influencing RBC storage quality and transfusion outcomes. Given caffeine’s short biological half-life, transient dietary modifications around the time of blood donation might mitigate its negative impact, aligning with blood donation guidelines in several European countries where donors are advised to limit caffeine intake prior to donation. Conversely, in other regions, such as the United States, caffeine consumption before blood donation is not actively discouraged and may even be implicitly encouraged due to its known beneficial acute effects on blood pressure, potentially expediting the donation process and reducing vasovagal reactions. Indeed, moderate caffeine intake can transiently increase donor blood pressure and vascular tone, facilitating venous access and blood withdrawal efficiency. However, this advantage must be balanced against caffeine’s mild diuretic properties, which may predispose donors to dehydration—an established risk-factor for adverse donation-related events and poorer blood flow during collection.

Beyond caffeine, our findings advocate for incorporating donor exposome data—dietary habits, medication usage—along with genetic markers into precision transfusion medicine strategies aimed at informing blood inventory management. For example, RBC units from high-caffeine-consuming donors or individuals with high-risk ADORA2b polymorphisms might be preferentially allocated to scenarios where slight variations in RBC quality are clinically less impactful. Conversely, units from donors with low caffeine exposure and favorable genetic profiles could be prioritized for high-risk transfusion recipients, such as neonates or patients with severe anemia.

Moreover, our mechanistic insights point to possible pharmacological interventions aimed at bolstering ADORA2b signaling or downstream metabolic pathways during RBC storage. Adenosine supplementation to standard storage additives transiently boosted energy metabolism, leading to early increases in intracellular adenosine and total adenylate pools, followed by elevation of purine deamination products inosine and IMP after storage week 3. Altogether, these results are consistent with adenosine uptake via ENT1, rather than activation of ADORA2b, followed by deamination of adenosine itself or its metabolite AMP into inosine and IMP by adenosine deaminase (27) or RBC-specific AMP deaminase 3 (43) Degradation of adenosine supplements in storage mediate would thus only favor transient stimulatory effects of ADORA2b, while the absence in stored RBC units of hepatic CYP2As that can degrade caffeine would not similarly constraint the antagonistic effects on the same receptors. In light of these results, alternative strategies to stimulate ADORA2b might include non-degradable adenosine analogs or allosteric modulators specifically enhancing glycolysis and redox balance, thereby improving stored RBC resilience and transfusion efficacy. Preliminary data from models utilizing AMPK activators (13) or hypoxic storage (25, 44) conditions provide proof-of-concept for these approaches, highlighting the feasibility of metabolic modulation as a storage enhancement strategy. Corollary to these considerations, cytochrome p450 reductases are NADPH dependent enzymes.(45) As such, direct G6PD inhibition by caffeine may constraint the pools of the very NADPH necessary for its direct catabolism or that of its metabolites like paraxanthine. Indeed, our mQTL analysis here identified an association between donor genetics (CYP2A6 SNPs, especially in donors of Asian and European descent) and paraxanthine metabolism, consistent with the literature (46, 47).

While our study benefits from robust epidemiological evidence combined with detailed mechanistic validation, we acknowledge certain limitations. The modest effect sizes observed in clinical transfusion outcomes raise questions about the absolute clinical relevance. Nevertheless, given the substantial global volume of transfusions, even small improvements in RBC quality can translate into meaningful clinical benefits at the population level. Additionally, our animal model, though highly informative, utilized defined genetic modifications; thus, translating these findings to routine clinical settings demands further validation.

Collectively, our study positions caffeine consumptionas a significant, modifiable factor influencing RBC metabolic responses to oxidant stress, here in the form of a common iatrogenic intervention such as storage under blood bank conditions. Our data thus suggests that caffeine might sensitize RBCs to hemolysis by osmotic or oxidant insults, common stressors for example in response to moderate (48) or strenuous exercise (49). It also underscores the importance of the gene-exposome interface in determining transfusion outcomes, advocating for precision approaches in transfusion medicine. Future investigations should further explore dietary guidance for donors, pharmacological enhancements of RBC metabolic pathways, and more comprehensive integration of donor metadata into clinical transfusion practices.

## Author Contribution

Animal studies: AH, JCZ. Proteomics: MD, KCH. Metabolomics and lipidomics analyses: DS, TN, AD. Biostatistics and Bioinformatics: GRK, XD, AD. REDS RBC Omics: MS, SK, SLS, PJN, MPB. Vein-to-vein database: NR; Human genetics: GRK, ALM, GPP (human). Figure preparation: AD. Conceptualization: YX, JCZ, AD. Writing and finalization: first draft by AD and all co-authors reviewed and approved the final version.

## Data and Materials availability

All raw data and elaborations are included in Supplementary Table 1.xlsx. ADORA2b KO mice are available upon reasonable request, finalization of material transfer agreement and after institutional ACUC approval through Dr James C Zimring Lab at the University of Virginia. Further information and requests for resources and reagents should be directed to and will be fulfilled by the Lead Contact, Angelo D’Alessandro.

## Data analysis and Statistical analyses

– including hierarchical clustering analysis (HCA), linear discriminant analysis (LDA), uniform Manifold Approximation and Projection (uMAP), correlation analyses and Lasso regression were performed using both MetaboAnalyst 5.0 and in-house developed code in RStudio (2024.12.1 Build 563).

## LIST of SUPPLEMENTARY MATERIALS

Supplementary Materials.pdf includes:

- SUPPLEMENTARY MATERIALS AND METHODS EXTENDED
- SUPPLEMENTARY FIGURES
- SUPPLEMENTARY FIGURE 1
- SUPPLEMENTARY FIGURE 2
- SUPPLEMENTARY FIGURE 3
- SUPPLEMENTARY FIGURE 4
- SUPPLEMENTARY FIGURE 5
- SUPPLEMENTARY FIGURE 6

Supplementary table 1.xlsx – including all raw data and elaborations.

