## Supplementary Methods and Figures for "Caffeine Impairs Red Blood Cell Storage Quality by Dual Inhibition of ADORA2b Signaling and G6PD Activity"

***Running title:*** *ADORA2b/G6PD Inhibition by Caffeine in stored RBCs*

#### Materials and Methods - Extended

##### RESOURCE AVAILABILITY

**Materials availability** ADORA2b KO mice are available upon reasonable request, finalization of material transfer agreement and after institutional ACUC approval through Dr James C Zimring Lab at the University of Virginia.

##### EXPERIMENTAL MODEL AND SUBJECT DETAILS

**Mouse RBC storage and post-transfusion recovery**

Mouse post-transfusion recovery (PTR) studies were performed as previously described (*1*). Storage of conditional, erythroid-specific ADORA2b KO mice (n=3) for 7 days was followed by transfusion into Ubi-GFP mice, which were used as recipients to allow visualization of the test cells in the non-fluorescent gate. To control for differences in transfusion and phlebotomy, Ubi-mCherry RBCs were used as a tracer RBC population (never stored). These RBCs were added to stored RBCs immediately prior to transfusion. PTR was calculated by dividing the post-transfusion ratio (Test/Tracer) by the pre-transfusion ratio (Test/Tracer), with the maximum value set as 1 (or 100% PTR).

**Donor recruitment in the REDS RBC Omics study:**

***Index donors*** A total of 13,758 donors were enrolled in the Recipient Epidemiology and Donor evaluation Study (REDS) RBC Omics at four different blood centers across the United States (https://biolincc.nhlbi.nih.gov/studies/reds_iii/). Of these, 97% (13,403) provided informed consent and 13,091 were available for metabolomics analyses in this study – henceforth referred to as "index donors". A subset of these donors were evaluable for hemolysis parameters, including spontaneous (n=12,753) and stress (oxidative and osmotic) hemolysis analysis (n=10,476 and 12,799, respectively) in ~42-day stored leukocyte-filtered packed RBCs derived from whole blood donations from this cohort (*2*). Methods for the determination of FDA-standard spontaneous (storage) hemolysis test, osmotic hemolysis (pink test) and oxidative hemolysis upon challenge with AAPH have been extensively described elsewhere (*3*).

***Recalled donors:*** A total of 643 donors scoring in the 5$^{th}$ and 95$^{th}$ percentile for hemolysis parameters at the index phase of the study were invited to donate a second unit of pRBCs, a cohort henceforth referred to as "recalled donors". These units were assayed at storage days 10, 23 and 42 for hemolytic parameters and mass spectrometry-based high-throughput metabolomics (*4*), proteomics (*5*), lipidomics (*6*) and ICP-MS analyses (*7*). Under the aegis of the REDS-IV-P project (*8*), a total of 1,929 samples (n=643, storage day 10, 23 and 42) were processed with this multi-omics workflow.

##### METHOD DETAILS

**High-throughput metabolomics** Metabolomics extraction and analyses in 96 well-plate format were performed as described, with identical protocols for human or murine RBCs(*9, 10*). RBC samples were transferred on ice on 96 well plate and frozen at -80 °C at Vitalant San Francisco (human RBCs) or University of Virginia (murine RBCs) prior to shipment in dry ice to the University of Colorado Anschutz Medical Campus. Plates were thawed on ice then a 10 uL aliquot was transferred with a multi-channel pipettor to 96-well extraction plates. A volume of 90 uL of ice cold 5:3:2 MeOH:MeCN:water (*v/v/v*) was added to each well, with an electronically-assisted cycle of sample mixing repeated three times. Extracts were transferred to 0.2 µm filter plates (Biotage) and insoluble material was removed under positive pressure using nitrogen applied via a 96-well plate manifold. Filtered extracts were transferred to an ultra-high-pressure liquid chromatography (UHPLC-MS —

Vanquish) equipped with a plate charger. A blank containing a mix of standards detailed before (*11*) and a quality control sample (the same across all plates) were injected 2 or 5 times each per plate, respectively, and used to monitor instrument performance throughout the analysis. Metabolites were resolved on a Phenomenex Kinetex C18 column (2.1 x 30 mm, 1.7 um) at 45 °C using a 1-minute ballistic gradient method in positive and negative ion modes (separate runs) over the scan range 65-975 m/z exactly as previously described.(*9*) The UHPLC was coupled online to a Q Exactive mass spectrometer (Thermo Fisher). The Q Exactive MS was operated in negative ion mode, scanning in Full MS mode (2 μscans) from 90 to 900 m/z at 70,000 resolution, with 4 kV spray voltage, 45 sheath gas, 15 auxiliary gas. Following data acquisition, .raw files were converted to .mzXML using RawConverter then metabolites assigned and peaks integrated using ElMaven (Elucidata) in conjunction with an in-house standard library(*12*).

**Lipid hydroperoxides and glutathionylated lipids: sample preparation:** Oxylipins were extracted via a modified protein crash from previously described.[10,14,16] Extraction of oxylipins from red blood cells were as follows: 10 μL of red cells was aliquoted directly into 90 μL of MeOH:MeCN:$H_2O$ (5:3:2, v:v:v) and pipetted up and down 5x per sample. Samples were then vortexed at 4 °C for 30 minutes. Following vortexing, samples were centrifuged at 12700 RPM for 10 minutes at 4 °C and 80 μL of supernatant was transferred to a new tube for analysis. 10 μL of extract from each sample was also combined to create a technical mixture, injected throughout the run for quality control. Lipidomics were extracted using identical prepartion techniques as oxylipins, save for utilizing MeOH:IPA (1:1, v:v) instead of MeOH:MeCN:$H_2O$ (5:3:2, v:v:v) as an extraction buffer.

**High-throughput Oxylipin Analysis:** Analyses were performed as previously published via a modified gradient optimized for the high-throughput analysis of oxylipins.[10,14,16] Briefly, the analytical platform employs a Vanquish UHPLC system (Thermo Fisher Scientific, San Jose, CA, USA) coupled online to a Q Exactive mass spectrometer (Thermo Fisher Scientific, San Jose, CA, USA). Lipid extracts were resolved over an ACQUITY UPLC BEH C18 column (2.1 x 100 mm, 1.7 μm particle size (Waters, MA, USA) using mobile phase (A) of 20:80:0.02 ACN:$H_2O$:FA and a mobile phase (B) 20:80:0.02 ACN:IPA:FA. For negative mode analysis the chromatographic the gradient was as follows: 0.35 mL/min flowrate for the entire run, 0% B at 0 min, 0% B at 0.5 min, 25%B at 1 min, 40%B at 2.5min, 55% B at 2.6min, 70% B at 4.5 min, 100% B at 4.6 min, 100% B at 6 min, 0% B at 6.1 min and 0% B at 7 min. The Q Exactive mass spectrometer (Thermo Fisher) was operated in negative ion mode, scanning in Full MS mode (2 μscans) from 150 to 1500 m/z at 70,000 resolution, with 4 kV spray voltage, 45 sheath gas, 15 auxiliary gas. Calibration was performed prior to analysis using the Pierce™ Positive and Negative Ion Calibration Solutions (Thermo Fisher Scientific).

**High-throughput Lipidomic Analysis:** Lipid extracts were analyzed (10 μL per injection) on a Thermo Vanquish UHPLC/Q Exactive MS system using a previously described(*6*) 5 min lipidomics gradient and a Kinetex C18 column (30 x 2.1 mm, 1.7 μm, Phenomenex) held at 50 °C. Mobile phase (A): 25:75 MeCN:$H_2O$ with 5 mM ammonium acetate; Mobile phase (B): 90:10 IPA:MeCN with 5 mM ammonium acetate. The gradient and flow rate were as follows: 0.3 mL/min of 10% B at 0 min, 0.3 mL/min of 95% B at 3 min, 0.3 mL/min of 95% B at 4.2 min, 0.45 mL/min 10% B at 4.3 min, 0.4 mL/min of 10% B at 4.9 min, and 0.3 mL/min of 10% B at 5 min. Samples were run in positive and negative ion modes (both ESI, separate runs) at 125 to 1500 m/z and 70,000 resolution, 4 kV spray voltage, 45 sheath gas, 25 auxiliary gas. The MS was run in data-dependent acquisition mode (ddMS$^2$) with top10 fragmentation. Raw MS data files were searched using LipidSearch v 5.0 (Thermo).

**Mass spectrometry-based Proteomics analyses:** Proteomics analyses were performed as described previously[78]. A volume of 10 μL of RBCs were lysed in 90 μL of distilled water containing 10 mM *N*-ethylmaleimide (NEM) for 20 min at room temperature. 5 μL of lysed RBCs were mixed with 45 μL of 5% SDS and then vortexed. Samples were reduced with 10 mM DTT at 55 °C for 30 min, cooled to room temperature, and then alkylated with 25 mM iodoacetamide in the dark for 30 min. Next, a final concentration of 1.2% phosphoric acid and then six volumes of binding buffer (90% methanol; 100 mM triethylammonium bicarbonate, TEAB; pH 7.1) were added to each sample. After gentle mixing, the protein solution was loaded to a S-Trap 96-well plate, spun at 1500 x g for 2 min, and the flow-through collected and reloaded onto the 96-well plate. This step was repeated three times, and then the 96-well plate was washed with 200 µL of binding buffer 3 times. Finally, 1 µg of sequencing-grade trypsin (Promega) and 125 µL of digestion buffer (50 mM TEAB) were added onto the filter and digested carried out at 37 °C for 6 h. To elute peptides, three stepwise buffers were applied, with 100 µL of each with one more repeat, including 50 mM TEAB, 0.2% formic acid (FA), and 50% acetonitrile and 0.2% FA. The peptide solutions were pooled, lyophilized, and resuspended in 500 µL of 0.1 % FA. Each sample was loaded onto individual Evotips for desalting and then washed with 200 µL 0.1% FA followed by the addition of 100 µL storage solvent (0.1% FA) to keep the Evotips wet until analysis. The Evosep One system (Evosep, Odense, Denmark) was used to separate peptides on a Pepsep column, (150 um inter diameter, 15 cm) packed with ReproSil C18 1.9 µm, 120A resin. The system was coupled to a timsTOF Pro mass spectrometer (Bruker Daltonics, Bremen, Germany) via a nano-electrospray ion source (Captive Spray, Bruker Daltonics). The mass spectrometer was operated in PASEF mode. The ramp time was set to 100 ms and 10 PASEF MS/MS scans per topN acquisition cycle were acquired. MS and MS/MS spectra were recorded from m/z 100 to 1700. The ion mobility was scanned from 0.7 to 1.50 $Vs/cm^2$. Precursors for data-dependent acquisition were isolated within ± 1 Th and fragmented with an ion mobility-dependent collision energy, which was linearly increased from 20 to 59 eV in positive mode. Low-abundance precursor ions with an intensity above a threshold of 500 counts but below a target value of 20000 counts were repeatedly scheduled and otherwise dynamically excluded for 0.4 min.

**Data analysis:** Acquired data was converted from raw to mzXML file format using Mass Matrix (Cleveland, OH, USA). Analysis was done using MAVEN, an open-source software program for oxylipin analysis. Lipid assignments and peak integration were performed using LipidSearch v 5.0 (Thermo Fisher Scientific). Samples were analyzed in randomized order with a technical mixture injected interspersed throughout the run to qualify instrument performance.

**Determination of hemoglobin and bilirubin increment via the vein-to-vein database** Association of ADORA2b SNPs – as a function of caffeine levels - with hemoglobin increments was performed by interrogating the vein-to-vein database, as described in Roubinian et al. (*13, 14*)

**Vein-to-vein database: General Study Design** We conducted a retrospective cohort study using electronic health records from the National Heart Lung and Blood Institute (NHLBI) Recipient Epidemiology and Donor Evaluation Study-III (REDS-III) program available as public use data through BioLINCC (*15, 16*). The database includes blood donor, component manufacturing, and patient data collected at 12 academic and community hospitals from four geographically diverse regions in the United States (Connecticut, Pennsylvania, Wisconsin, and California) for the 4-year period from January 1, 2013 to December 31, 2016. Genotype and metabolomic data from the subset of blood donors who participated in the REDS-III RBC-Omics study (*17*) was linked to the dataset using unique donor identifiers.

**Study Population and Definitions** Available donor genetic polymorphisms for ADORA2b were linked to issued RBC units using random donor identification numbers. Among transfusion recipients, we included all adult patients who received a single RBC unit during one or more transfusion episodes between January 1, 2013 and December 30, 2016. Recipient details included pRBC storage age, and blood product issue date and time. We collected hemoglobin levels measured by the corresponding clinical laboratory prior to and following each RBC transfusion event (0h and 24h after transfusion).

**Transfusion Exposures and Outcome Measures** All single RBC unit transfusion episodes linked to genetic polymorphism for ADORA2b were included in this analysis. A RBC unit transfusion episode was defined as any single RBC transfusion from a single donor with both informative pre- and post-transfusion laboratory measures and without any other RBC units transfused in the following 24-hour time period. The outcome measures of interest were change in hemoglobin (ΔHb; g/dL) following a single RBC unit transfusion episode. These outcomes were defined as the difference between the post-transfusion and pre-transfusion levels. Pre-transfusion thresholds for these measures were chosen to limit patient confounding (e.g., underlying hepatic disease). For pre-transfusion hemoglobin, the value used was the most proximal hemoglobin measurements prior to RBC transfusion, but at most 24 hours prior to transfusion. Furthermore, we excluded transfusion episodes where the pre-transfusion hemoglobin was greater than 9.5 g/dL, and the hemoglobin increment may be confounded by hemorrhage events. For post-transfusion hemoglobin, the laboratory measure nearest to 24-hours post-transfusion, but between 12- and 36-hours following transfusion was used.

##### Storage of human packed RBCs in presence of adenosine

Leukocyte filtered packed RBCs (n=3 in experiment 1 and n=5 in experiment 2) were stored in standard CP2D-AS-3 (additive solution 3), either untreated or supplemented with 1, 5 or 10 µM adenosine (no. A9251- Sigma Aldrich; experiment 1) or 10 µM (experiment 2), a concentration below 14 µM – which would potentially be arrhythmia-inducing in transfusion recipients via stimulation of adenosine 1 receptor in cardiac atria (*18*).

#### QUANTIFICATION AND STATISTICAL ANALYSIS

**Statistical Methods** We assessed the univariable association of all *a priori* selected donor, manufacturing, and recipient covariates with the outcomes using linear regression. Multivariable linear regression assessed associations between alleles of donor single nucleotide polymorphisms and changes in hemoglobin levels post-transfusion hemoglobin. Two-sided p-values less than 0.05 were considered to be statistically significant. Analyses were performed using Stata Version 14.1, StataCorp, College Station, TX.

**Data analysis** Statistical analyses – including hierarchical clustering analysis (HCA), linear discriminant analysis (LDA), uniform Manifold Approximation and Projection (uMAP), correlation analyses and Lasso regression were performed using both MetaboAnalyst 5.0 (*19*) and in-house developed code in RStudio (2024.12.1 Build 563).

**REDS RBC Omics mQTL analyses of paraxanthine:** The workflow for the mQTL analysis of paraxanthine and theobromine levels in REDS Index donors is consistent with previously described methods from our pilot mQTL study on 250 recalled donors (*20*). Details of the genotyping and imputation of the RBC Omics study participants have been previously described by Page, et al. (*21*) Briefly, genotyping was performed using a Transfusion Medicine microarray (*22*) consisting of 879,000 single nucleotide polymorphisms (SNPs); the data are available in dbGAP accession number phs001955.v1.p1. Imputation was performed using 811,782 SNPs that passed quality control. After phasing using Shape-IT (*23*), imputation was performed using Impute2 (*24*) with the 1000 Genomes Project phase 3 (*24*) all-ancestry reference haplotypes. We used the R package SNPRelate (*25*) to calculate principal components (PCs) of ancestry. We performed association analyses for paraxanthine using an additive SNP model in the R package ProbABEL (*26*) and 13,091 REDS Index donor study participants who had both paraxanthine and theobromine levels measured. We adjusted for sex, age (continuous), frequency of blood donation in the last two years (continuous), blood donor center, and ten ancestry PCs. Statistical significance was determined using a p-value threshold of $5x10^{-8}$. We only considered variants with a minimum minor allele frequency of 1% and a minimum imputation quality score of 0.80. The OASIS: Omics Analysis, Search & Information a TOPMED funded resources (*27*), was used to annotate the top SNPs. OASIS annotation includes information on position, chromosome, allele frequencies, closest gene, type of variant, position relative to closest gene model, if predicted to functionally consequential, tissues specific gene expression, and other information.

### SUPPLEMENTARY FIGURES

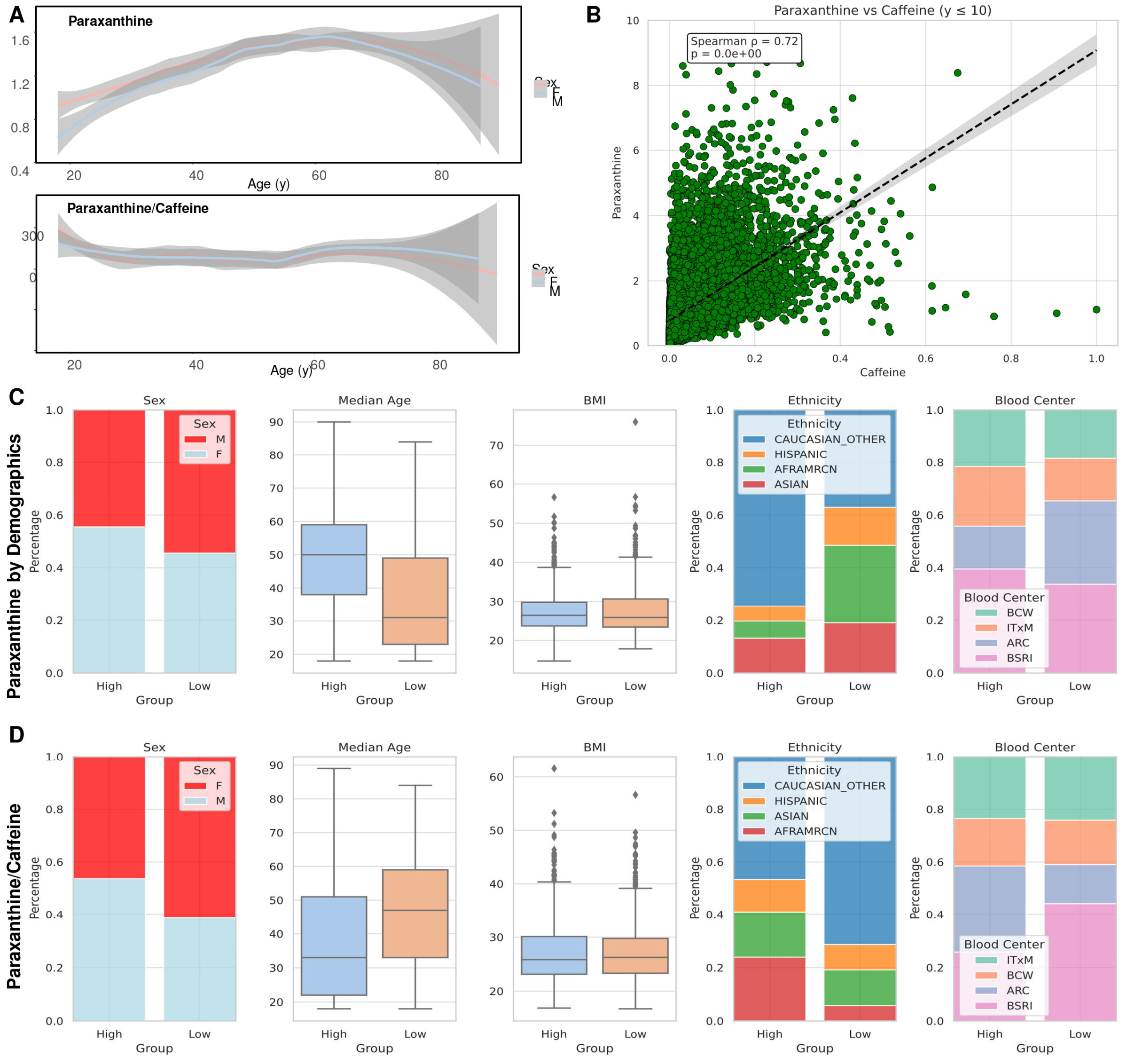

**Supplementary Figure 1** –In **A**, paraxanthine levels in 13,091 Index REDS RBC Omics blood units as a function of donor age and sex. In **B**, significant correlation between caffeine and paraxanthine levels. Stratification of paraxanthine (**C**) and paraxanthine/caffeine ratios (**D**) by donor sex, age, BMI, self-reported ethnicity and blood center in which the donors were enrolled.

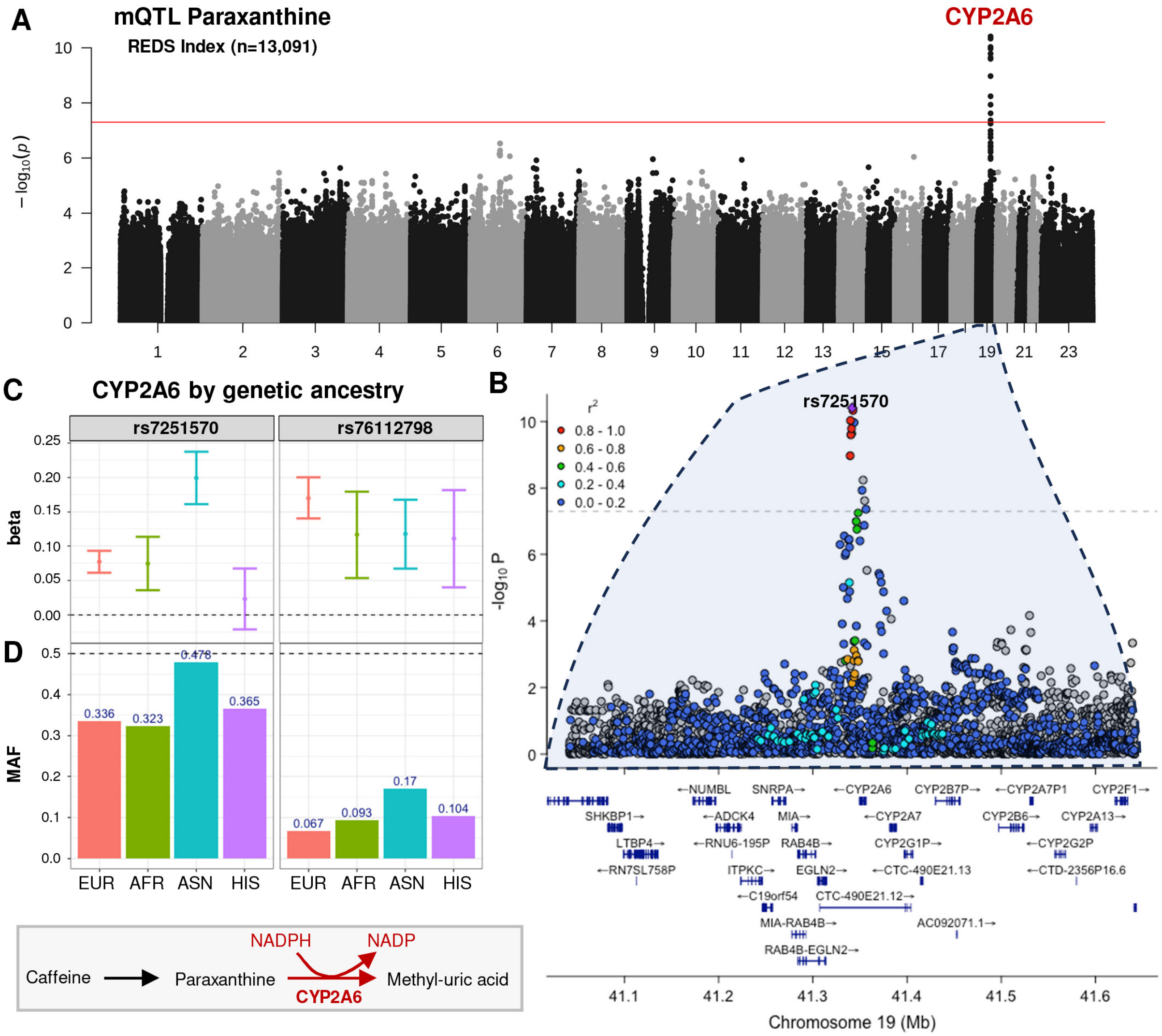

**Supplementary Figure 2** –Metabolite Quantitative Trait Loci (mQTL) analysis for paraxanthine identified a region on chromosome 19 significantly associated with heterogeneity in the level of this metabolite in the 13,091 REDS Index packed RBC units (**A**). Locus Zoom analysis for the total population (**B**) and separate analysis by donor genetic ancestry identified three key single nucleotide polymorphisms (SNPs) in the region coding for CYP2A6, rs7251570, rs76112798 and rs11667314 as the top hits for the total donor populations, or for donors of Asian or European descent, respectively (**C-D**), with rs11667314 and rs721570 likely linked and part of the same haplotype.

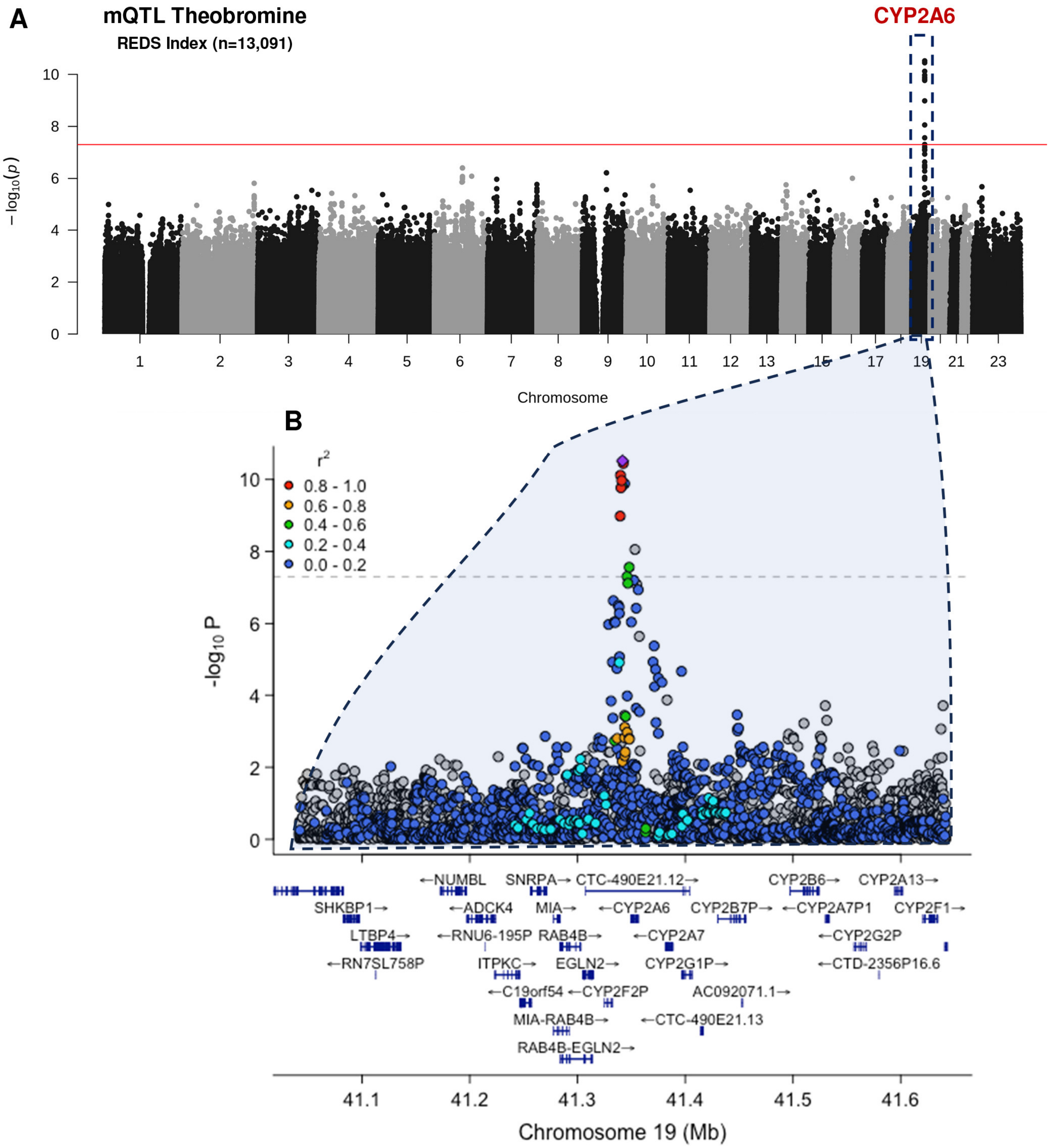

**Supplementary Figure 3** –Metabolite Quantitative Trait Loci (mQTL) analysis for theobromine identified a region on chromosome 19 significantly associated with heterogeneity in the level of this metabolite in the 13,091 REDS Index packed RBC units (**A**). Locus Zoom analysis for the total population (**B**).

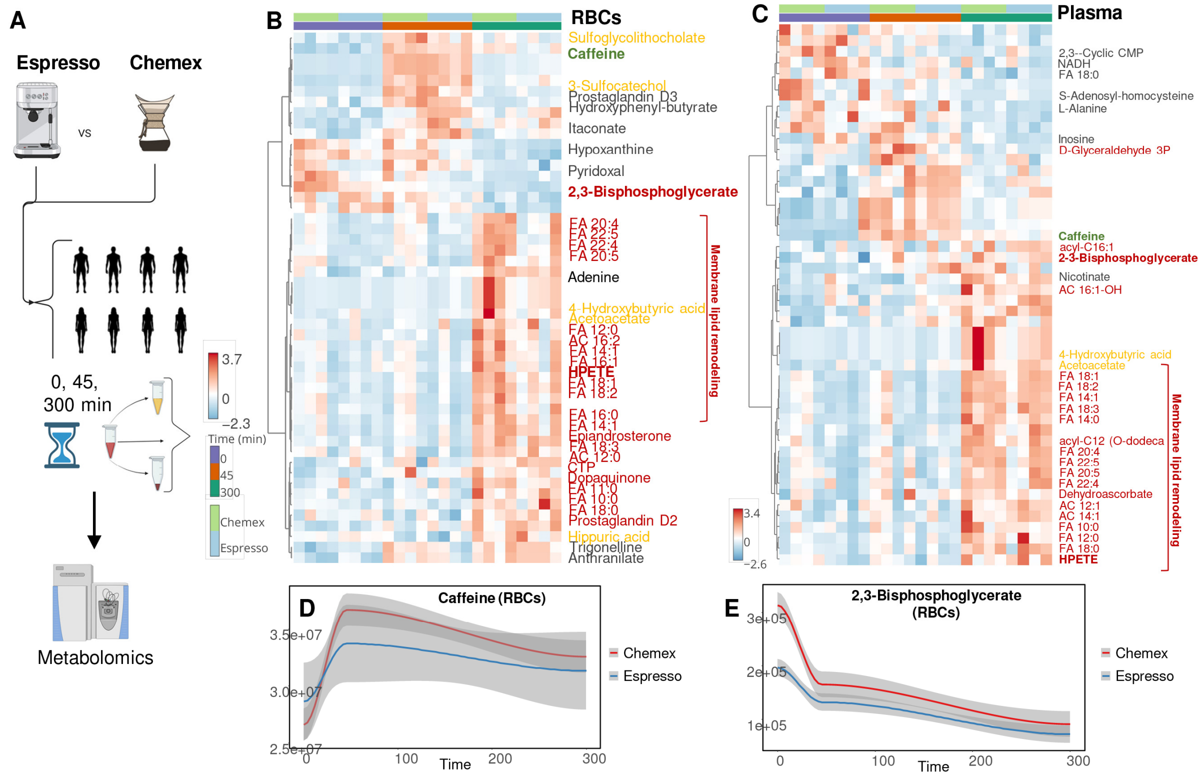

**Supplementary Figure 4** –Eight volunteers were randomized to consume a cup of coffee brewed either via Chemex or an espresso, prior to blood collection for plasma and RBC metabolic characterization at baseline, after 45 min or 5h (300 min) after consuming the beverage (**A**). In **B-C**, hierarchical clustering analysis of the top 50 metabolites in RBCs and plasma by linear discriminant analysis as a function of time and brewing method. Metabolites highlighted include caffeine (green), bacterial metabolites (orange), glycolytic metabolites, free fatty acids, acyl-carnitines and lipid oxidation products as a marker of membrane lipid remodeling (dark red). In **D-E**, line plots for RBC caffeine and 2,3-bisphosphoglycerate as a function of time and brewing method.

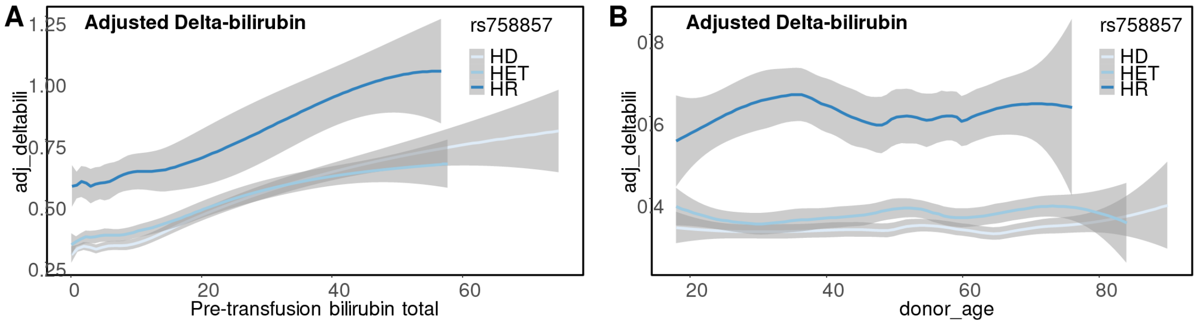

**Supplementary Figure 5** – Delta bilirubin levels (adjusted by donor/recipient weight, sex – y axis) by pre-transfusion bilirubin levels (**A**) or donor age (**B**) as a function of rs785587 SNPs (0 = homozygote dominant; 1 = heterozygote; 2 = homozygote for the alternative or recessive allele – HR).

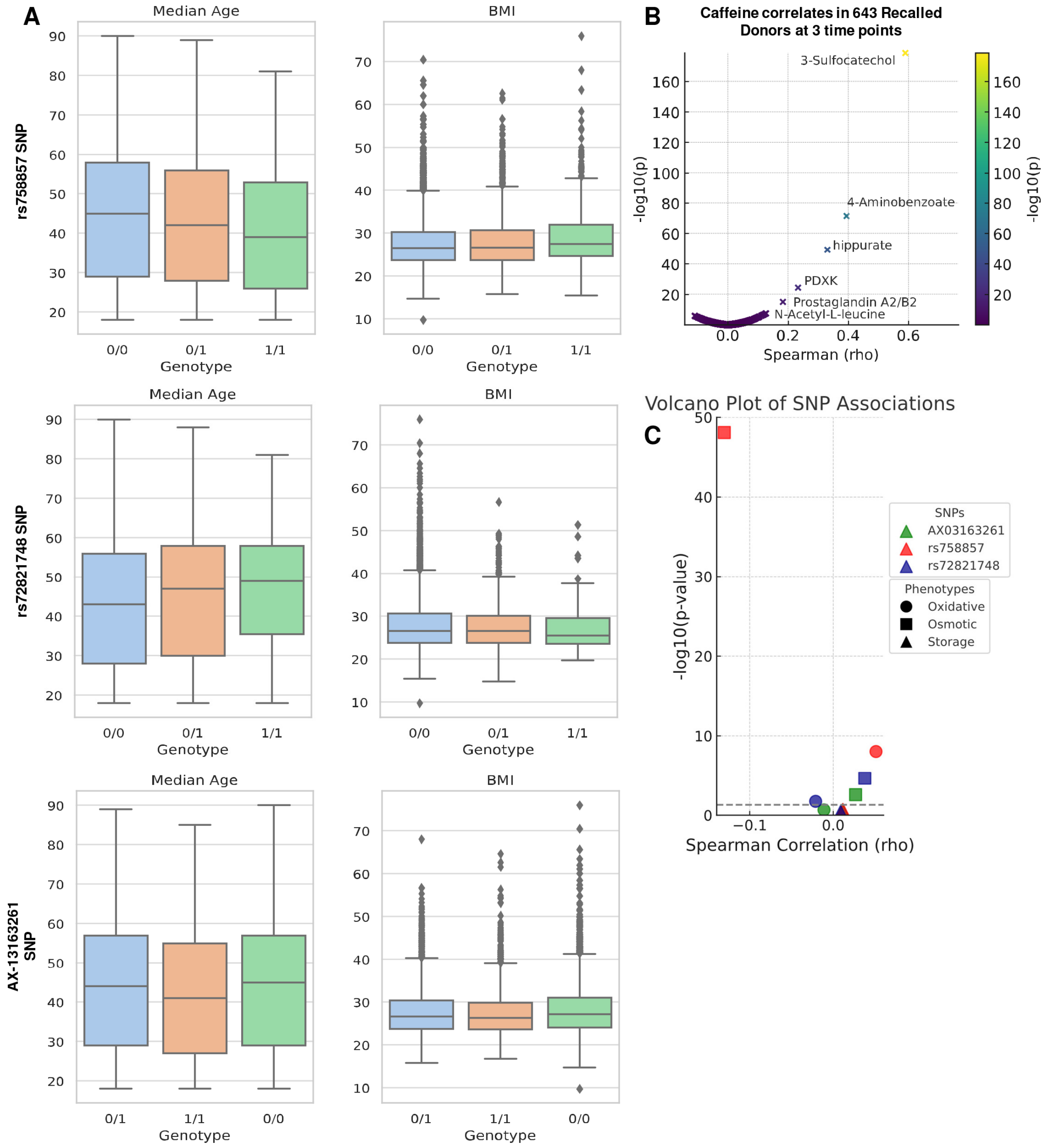

**Supplementary Figure 6** – Stratification of ADORA2b most prevalent SNPs as a function of donor age and BMI in the 13,091 REDS RBC Omics Index cohort (**A**). In **B**, correlates of caffeine levels to metabolites in the 643 REDS RBC Omics Recalled donors (**B**). In **C**, volcano plot of Spearman correlation (x axis) and significance (-log10 of p-values – y axis) of ADORA2b most prevalent SNPs and hemolysis parameters.

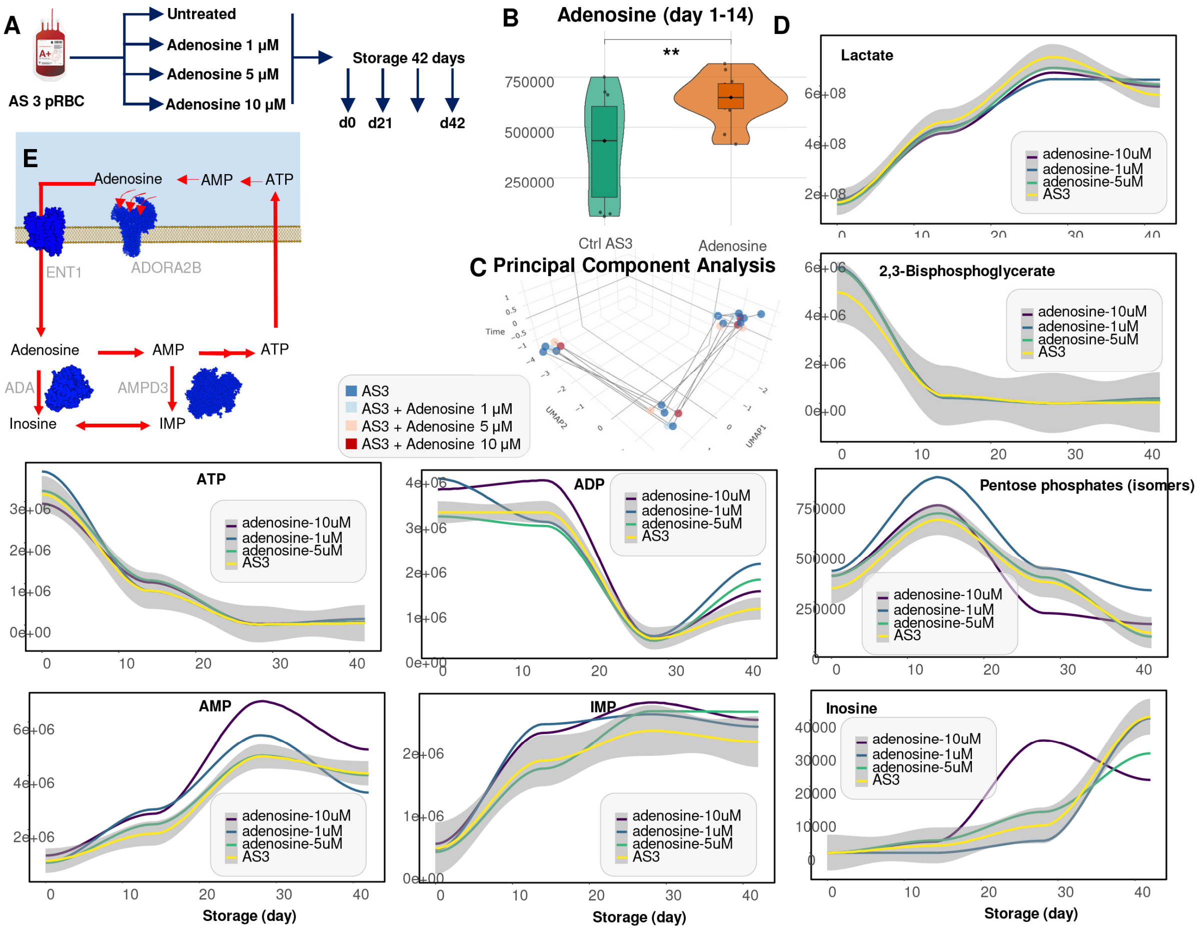

**Supplementary Figure 7** – Storage of human packed RBCs in presence of adenosine (**A**) did increase intracellular level of this metabolite during the first 2 weeks of storage (**B**), but had minimal effect on the metabolome – which overlapped between controls and supplemented cells (**C**). Changes were noted in glycolysis (lower levels of lactate – line plots in **D**), 2,3-bisphosphoglycerate and ATP or ADP (first week of storage), and AMP, IMP and inosine after storage week 3. Altogether, these results are consistent with adenosine uptake via ENT1, rather than activation of ADORA2b, followed by deamination of adenosine itself or its metabolite AMP into inosine and IMP by adenosine deaminase or RBC-specific AMP deaminase 3 (**E**).
